# Age-associated DNA methylation changes in *Xenopus* frogs

**DOI:** 10.1101/2022.07.22.500491

**Authors:** Marco Morselli, Ronan Bennett, Nikko-Ideen Shaidani, Marko Horb, Leonid Peshkin, Matteo Pellegrini

**Affiliations:** Molecular, Cell & Developmental Biology, UCLA, Los Angeles, CA 90095, USA; Systems Biology, Harvard Medical School, Boston, MA 02115, USA; Eugene Bell Center for Regenerative Biology and Tissue Engineering and National Xenopus Resource, Marine Biological Laboratory, Woods Hole, MA 02543, USA

**Author notes:** Department of Chemistry, Life Sciences and Environmental Sustainability, University of Parma, Parma, 43124 Italy (Marco Morselli).

## Abstract

Age-associated changes in DNA methylation have been widely established and characterized in mammals but not yet extended to other species. We present clear evidence that the aquatic vertebrate species *Xenopus tropicalis* displays clear patterns of age-associated changes in DNA methylation. This observation allows us to leverage the unique resources available for *Xenopus* to study how DNA methylation relates to other hallmarks of aging. We have generated whole genome bisulfite sequencing (WGBS) profiles in nine samples representing young, mature and old adults and characterize the gene- and chromosome-scale changes. Moreover, we selected a few thousands CpG sites to expand the findings in a larger cohort of individuals through targeted DNA methylation analysis to produce a cost effective epigenetic clock assay.

## Introduction

Amphibians have not been extensively examined in aging studies, yet *Xenopus* has many unique features as a prospective model organism for aging biology. It is thought to have negligible senescence [Brooks’61, Kara’94], remains fertile late in life, provides an ease of surgical and biochemical manipulation of the embryos and genome editing, and large oocytes. To date, it could not be used as an aging model due to the lack of verifiably old animals and a long lifespan (estimated maximum lifespan of over 30 years in *X.laevis* and 12 years in shorter lived *X. tropicalis*). If methylation clocks worked in *Xenopus* it would enable a wide range of new inquiries. The 2012 Nobel Prize in Medicine was awarded to Sir John Gurdon for his work on nuclear transfer and reprogramming in *Xenopus*, which he chose for the ease of injection of somatic nuclei into oocytes. This well established model system was used for numerous discoveries in biology over many decades. To date *Xenopus* has not been considered a model organism for aging research even though some research into reproductive system aging in *Xenopus* has been carried out [Brooks et al., 1961, Kara et al., 1994].

There are certain characteristics of aging in amphibians which make them extremely attractive aging models, such as old age maintenance of reproductive performance, neurogenesis, myogenesis, and teeth replacement in some species of amphibians. Chiefly used as a model for embryology, *Xenopus* adult females are often culled as early as 5 y/o since young healthy adults are easily available. Males are routinely sacrificed to procure testes. Establishing *Xenopus* as an aging model requires collecting a cohort covering a wide range of confidently established ages, followed by validating and calibrating a methylation clock. To date methylation clocks have been largely limited to mammalian species. There are papers showing variable methylation with age in some non-mammalian species but these studies stop short of calibrating robust clocks.

The National Xenopus Resource at the Marine Biology Lab has been maintaining the inbred *Xenopus* animals for over 10 years creating a unique collection of animals of known age, covering ages from 1 to 10 years. We have established protocols for sampling the skin without sacrificing the animal.

It has been established that using WGBS (whole-genome bisulfite sequencing) CpG sites show an average 91% methylation over different developmental stages in *X. tropicalis* [Bogdanović’ et al., 2011; Bogdanović’ et al., 2016], including gametes (sperm and spermatids) [Teperek, et al. 2016]. Previous reports have shown that DNA methylation and histone modifications are highly interconnected in *X. tropicalis*, and similarly to mammals, 5^me^C levels are anticorrelated with H3K4me3 and H3K27me3 marks [Bogdanović’ et al., 2011; Hontelez et al., 2015; Session et al., 2016], with many promoters being hypomethylated [Long et al., 2013]. Xenopus genomes contain two DNA cytosine (C5) DNA methyltransferases (DNMTs): DNMT1 and DNMT3a, a maintenance and a *de novo* DNA methyltransferase, respectively [Iwasaki Y, et al., 2014; Kyono, et al, 2016] [Supplementary Figure 1]. In mammals, in addition to the previously mentioned DNMTs, there are an additional de novo enzyme (DNMT3b), an inactive DNMT-like accessory factor (DNMT3L), and a rodent-specific enzyme involved in male germ line retrotransposon silencing (DNMT3c) [Jurkowska and Jeltsch, 2016; Schmitz et al., 2019].

Using established cohorts for cross-sectional study of DNA methylation, we sampled skin patches of nine animals, to evaluate the feasibility of creating methylation clocks in aging *Xenopus tropicalis*.

## Results

In order to assess the age-related changes, we chose to focus on three ages, covering the range of ages from young to old adults. The National Xenopus Resource has maintained the animal stocks for 10 years at the point of the sampling thus the oldest animals represent some of the oldest available individuals known to the Worldwide Xenopus research community. The youngest animals represent adults which have been reproducing for a few months as sexual maturity in *X. tropicalis* comes at 4-6 months depending on husbandry conditions [McNamara et al, 2018 and Shaidani et al 2020].

Cytosine DNA methylation was measured by WGBS (see Materials and Methods). The samples were collected from the skin of the webbing in the hindlimb of nine individuals of *Xenopus tropicalis*. The resulting methylation showed a distribution typical of the methylation of higher eukaryotes (Fig. 1A), with the majority of the CpG sites fully methylated or highly methylated (>80%), and only a small fraction completely unmethylated. A similar pattern is observed in mammals as exemplified by the human mammary epithelial tissue (Fig 1A). DNA methylation in other contexts (CpHpH and CpHpG) was extremely low (Table 1). The small differences in the patterns between the frog and human samples could likely be attributed to the differences in tissue purity w.r.t. A cell type composition between the epithelial samples in human and skin punch in frog.

**Figure 1:**
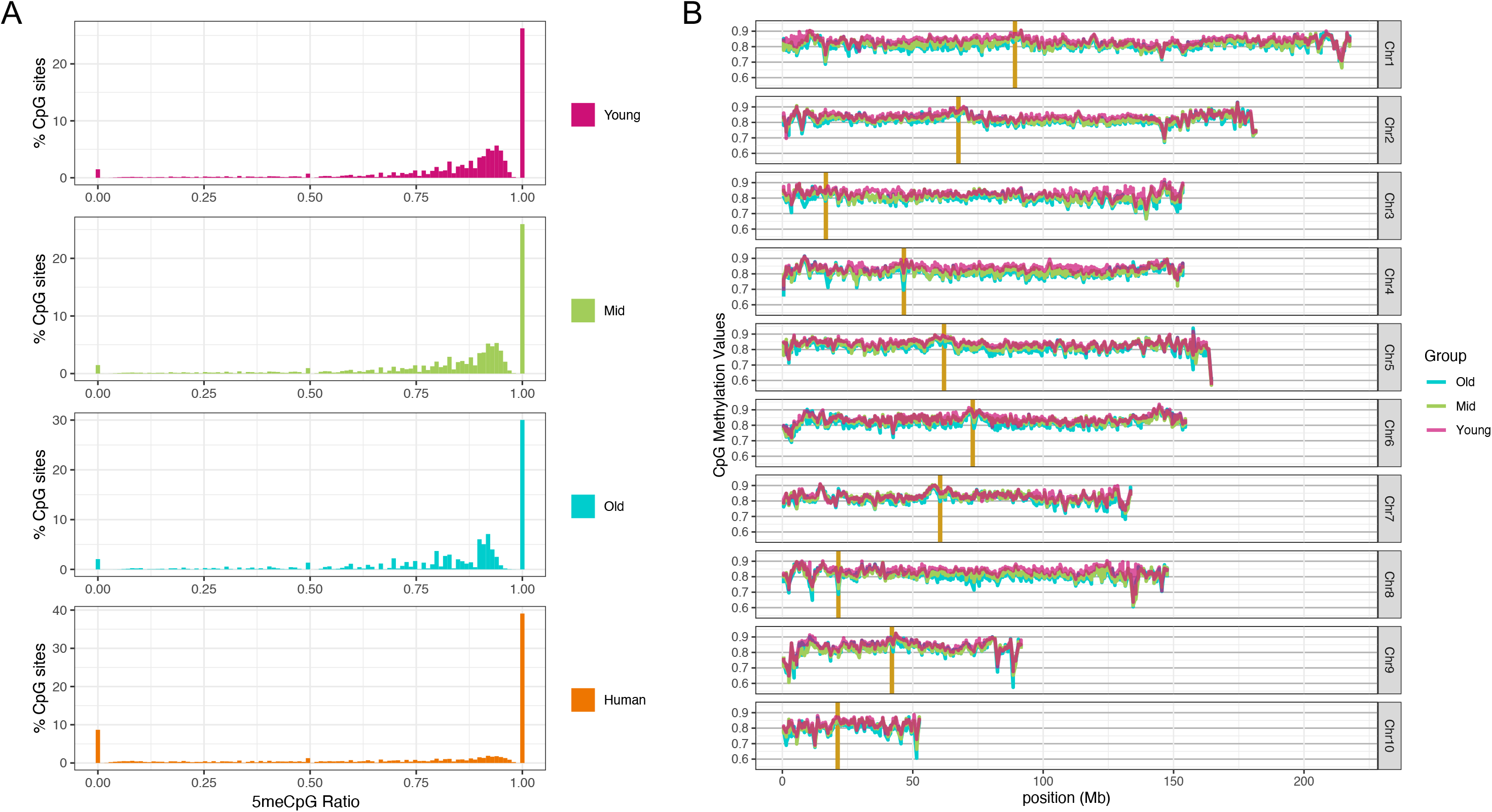
Cytosine methylation levels in CpG context. (A) Distribution for X. tropicalis of three different age groups and human mammary epithelial tissue. For each X. tropicalis age group the average of three different individuals. Common sites with at least 5x coverage were used. The human mammary epithelial dataset is from ENCODE (doi:10.17989/ENCSR656TQD, file ENCFF699GKH). (B) CpG methylation distribution over chromosomes for each age group (average of all samples within the same group). Vertical gold bars represents centromere positions. Common sites to all samples with at least 3x coverage were used. Resolution 500Kb.

**Table 1:** Average 5^me^C (%) by dinucleotide context

| *Sample* | *Age (yrs)* | *CpG* | *CpA* | *CpC* | *CpT* |
| --- | --- | --- | --- | --- | --- |
| Young_1 | 1 | 81.83 | 00.93 | 00.54 | 00.59 |
| Young_2 | 1 | 82.17 | 00.93 | 00.56 | 00.61 |
| Young_3 | 1 | 81.88 | 00.84 | 00.51 | 00.53 |
| Mid_1 | 5 | 79.82 | 00.83 | 00.48 | 00.50 |
| Mid_2 | 5 | 80.90 | 00.83 | 00.50 | 00.53 |
| Mid_3 | 5 | 79.80 | 00.86 | 00.53 | 00.55 |
| Old_1 | 9 | 78.69 | 01.38 | 00.74 | 00.95 |
| Old_2 | 9 | 79.75 | 00.99 | 00.56 | 00.62 |
| Old_3 | 9 | 79.93 | 00.93 | 00.54 | 00.58 |

Taking advantage of recently published high quality chromosome-level assembly of the *X.tropicalis* genome, we examined chromosome-level DNA methylation patterns (Fig. 1B). The CpG methylation levels over 500 Kb-bins are high across the genome with only a few dips towards the ends of a few chromosomes (e.g. subtelomeric regions of Chr5, and Chr9), but never below 60%. At his level of signal smoothing, there is a noticeable difference in how uniform the levels of DNA methylation are across the chromosomes - towards the telomeric regions, there is much higher variability in every chromosome. This spatial pattern is conserved across all three ages. Since pero-centromeric chromatin is known to be highly repetitive, and DNA methylation patterns are known to decorate the genomic repeats, we noted the centromeres of the *X.tropicalis* chromosomes. The centromeric regions of most of the chromosomes (Chr 2, 4, 6, 7, 8 and 9) appear to be hypo-methylated similarly to the human DNA (Gershman et al 2022).

Consistent with what we know from mammalian DNA methylation patterns, the global methylation levels of CpG sites are slightly but statistically significantly different among the three age groups. The highest levels are seen in the ‘young’ group (approx. 1 y/o), intermediate in ‘Mid’ (approx. 5 y/o), and the lowest in the Old’ group (approx. 9 y/o) (Table 1). The same pattern of methylation reduction with age is clearly seen in both genes (Fig 2A) and repeated elements (Fig. 2B). Similarly to mammalian 5^me^C distribution, the gene body is highly methylated (average methylation levels are on above 75%) and the region within 1Kb of the TSS show lower levels of DNA methylation, as previously reported [Long et al., 2013]. This can be due to the presence of histone marks repulsive for DNA methylation (e.g. H3K4 methylation) for expressed genes, or other histone-based repressive marks (e.g. H3K27 methylation) for non expressed genes. Conversely, repeated elements (longer than 1 Kb) show high methylation levels (average >80%) spanning their entire length. The methylation levels of common CpG sites can also be used to discriminate the samples of the three age groups through a principal components analysis (Fig. 2C). Principal Component 1 (PC1) has a nearly perfect correlation with age (R^2^ = 0.98), while other PCs show no correlation. Here again consistent with mammalian age-related methylation patterns we see a much clearer concordance across the young samples compared to the old samples with middle age samples demonstrating intermediate variability.

**Figure 2:**
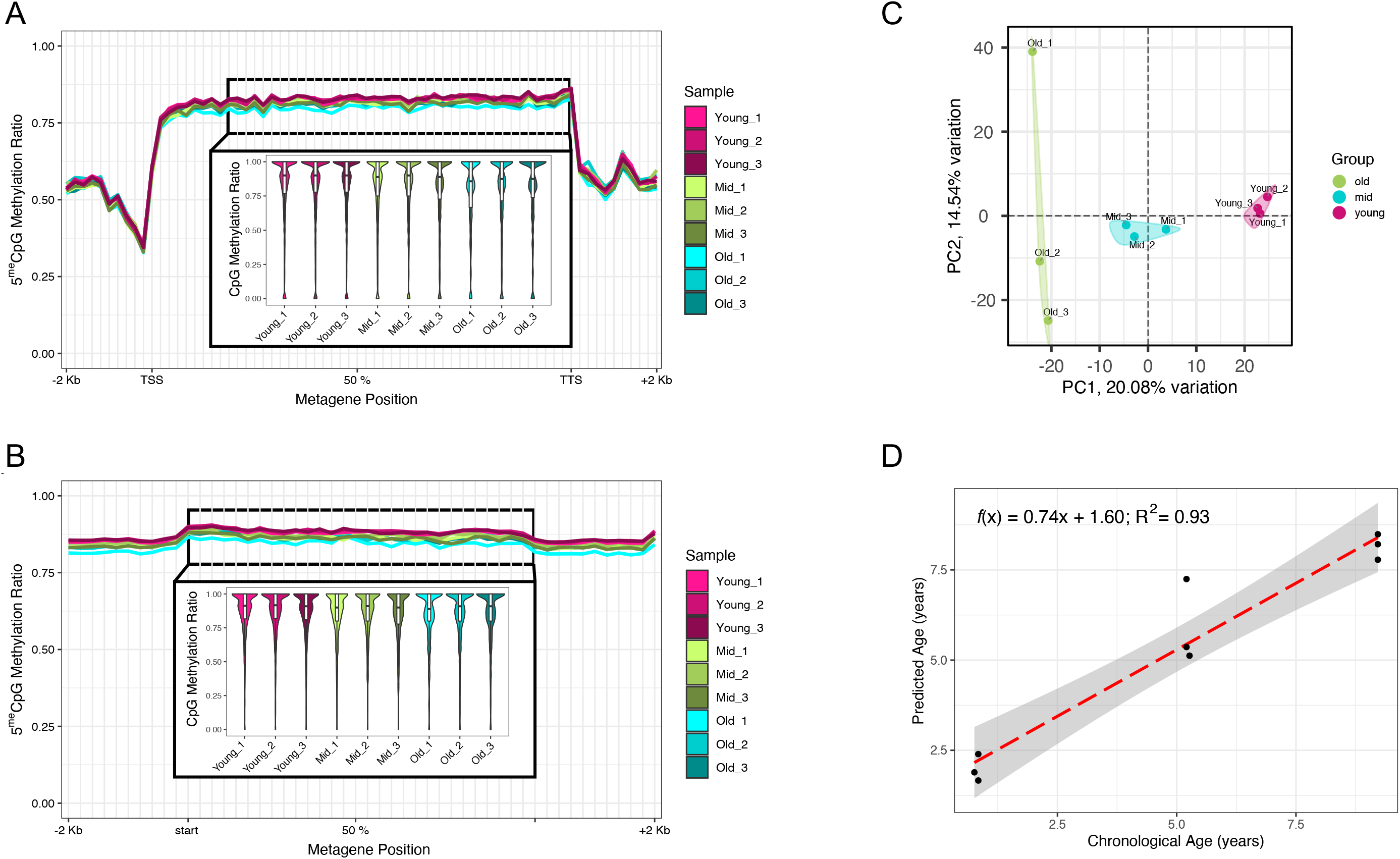
CpG methylation distribution over genomic features. (A) gene metaplot with upstream and downstream 2Kb. In the inset, a zoom in of the central part of the gene body (from the second decile to the TTS). (B) Gene body CpG methylation in repeated elements bigger than 1Kb [not distinguished by class nor family]. (C) Principal Component Analysis (showing PC1 vs. PC2) using CpG methylation levels for the nine frog samples. (D) CpG methylation clock leave-one-out predictions. Each point represents the age prediction on one sample from an Elastic Net model which was trained on the other 8 samples. For all panels, common sites to all samples with at least 5x coverage were used. The boxplots in the insets show the mean of the distribution.

Encouraged by such clear differences, we next asked whether we can use the CpG methylation levels to predict the age of an individual. Normally a much larger number of samples would be required to build and validate a methylation-based aging model, but three ages with three repeats each already allow us to construct a preliminary clock. We used Elastic Net regularized linear regression for the model. The predictors were the methylation values at all CpG sites across the genome with 5x coverage (110544 sites). By performing leave-one-out cross validation (LOOCV), we achieved an excellent correlation between actual age and predicted age (R^2^ = 0.93) (Fig. 2D). However, the age predictions are a slight overestimate for the young frogs, and an underestimate for the old frogs. The LOOCV procedure involved fitting 9 Elastic Net clock models, which altogether selected 331 distinct CpG sites as relevant features (Supplementary Figure 2).

In order to expand and confirm the findings in a bigger cohort, while avoiding the high costs of WGBS for a large genome, we selected a few thousand sites (approx. 3000) that are strongly (both positively and negatively) correlated with age (Fig. 3A). Using the CpG methylation levels of the selected sites, we were able to cluster the nine samples based on age groups (Fig. 3B). As expected, the majority of the sites show a noticeable decrease in DNA methylation levels from young to old samples (green and purple clusters), but a fraction (approx. 10% - gold cluster) shows an inverse pattern. Notice that there are hundreds of highly informative sites with different age associated patterns. Some sites are modified between young and middle ages, while others between the middle and old. It is clear that while overall similarity of the DNA methylation patterns is diminishing with age, the most informative sites are most discordant in the middle age, suggesting that the trajectories from young to old diverge in the middle age. The selected sites are mainly found within genes and repeated elements, and only a small percentage can be found outside of the elements, but always within 1Kb (Supplementary Figure 3).

**Figure 3:**
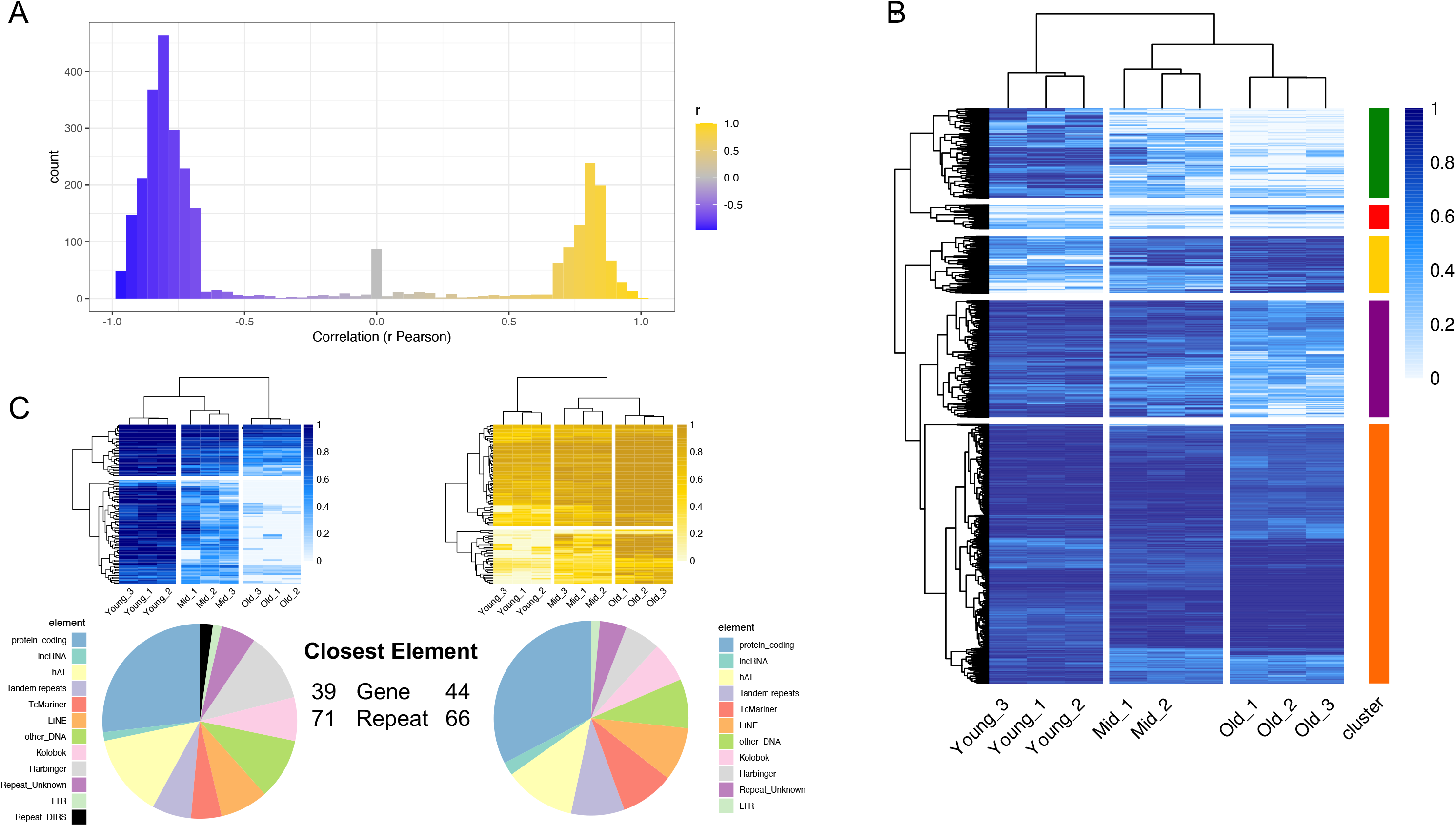
Selection of CpG sites correlated with age. (A) Pearson correlation (r) distribution of the approximately 3000 CpG sites selected. (B) CpG methylation levels of the selected sites (same as in A). Sites are grouped in 5 clusters, based on the clustering (method = ‘complete’). (C) CpG methylation levels of the top 100 sites correlated with age (right, gold), and anticorrelated (left, blue) (top). Distribution of the closest genomic elements (both genes and repeats) for each site (bottom).

## Conclusions and Discussion

We have characterized the age-associated changes in DNA methylation in *Xenopus tropicalis* frogs of three different ages covering the span of young, mature and old adult ages. Our findings indicate that DNA methylation clocks will be reliable and robust tools of investigation of aging biology outside of the mammalian species, and to the best of our knowledge the *X. tropicalis* clock will be the first amphibian methylation clocks. Many of the methylation features and changes we observe are consistent with what is known in mammalian species, suggesting that the mechanism of age-related changes are conserved. The sites selected in this study will be used to design probes and perform targeted methylation through capture hybridization in a larger cohort of samples.

The present work is based on skin samples which are easily available and do minimal harm to the animals, with repeated sampling possible after animals are given a few months break, which allows one to study longitudinal changes in one animal. We have not attributed specific methylation changes to specific cell types, even though a webbing sample contains not only skin but also connective tissue and blood. We expect that methylation clocks will be applicable to distinct frog tissues just as they are in mammals. One strength of Xenopus as a model organism is in embryology. Since the embryo is developing outside of the mother’s organism, it is easily accessible for manipulation and observation, including embryonic grafts from transgenic animal lines. This in turn opens up new directions for studying the causality of methylation and tracking the establishment of methylation patterns for the sites, correlated with age across cell types.

Crucially, being able to estimate the age of an individual frog opens up an exciting possibility of obtaining samples from wild animals. As *Xenopus* is relatively long lived and seldom kept in animal facilities past 3 years, the ability to cheaply attribute age to the animal with a targeted assay which is likely to generalize across *Xenopus* species is an important step in understanding the causal role of DNA methylation is age-related disease and organ malfunction.

## Materials and Methods

### Sample collection and DNA extraction

Nine sampled adult *Xenopus tropicalis* were housed at the National Xenopus Resource (RRID:SCR_013731) in two multi-rack recirculating aquatic systems with established diet and water parameters (conductivity, pH, and temperature) as previously defined [McNamara et al, 2018 and Shaidani ett al, 2020].

Disposable biopsy punches were used to collect tissue (VWR 21909–140) from the hindlimb webbing of mutant, transgenic, and wild type *X. tropicalis* with Nigerian St.549 background (RRID_NXR_1018).

DNA extraction was performed from web punches (4 mm in diameter except 6months old for which we did 2mm), according to Xu Y. et al. (Xu et al. 2020) with minor modifications. Briefly, tissue chunks were digested o/n in a 1.5 ml tube containing 200 μl of lysis buffer (100 mM Tris-HCl pH 8; 200 mM NaCl; 0.20% SDS; 5 mM EDTA) + 4 μl of Proteinase K (20 mg/ml, NEB) at 55°C (300 rpm continuous shaking in a thermomixer). After centrifugation for 15 min at 16’000g (room temperature), the supernatant was transferred into a new tube avoiding the debris. The centrifugation step was repeated after the addition and mixing of the same volume of isopropanol. The resulting pellet was then washed twice with 500 μl of EtOH 70% (centrifugation at 16’000g for 10 min, RT). The remaining ethanol was then carefully removed and the pellet air dried for 5-10 minutes at 55°C (open caps). The dried pellet was then resuspended with 55 μl of EB buffer (10 mM Tris-HCl pH 8) at 55°C for 1 h at 1400 rpm in a thermomixer, before being quantified (Qubit dsDNA BR - LifeTechnologies) and quality checked (Agilent 4200 TapeStation - Genomic Assay).

### WGBS Library Preparation

1 μg of purified DNA was sonicated using the Bioruptor Pico (Diagenode) for 15 cycles of 30 second ON / 90 seconds OFF. Libraries were prepared according to Morselli et al. (Morselli et al. 2022)with minor modifications. Briefly, NEB Next Ultra II DNA kit was used for end-repair, A-tailing and ligation of pre-methylated unique-dual indexed adapters (Morselli et al. 2021). Bisulfite conversion was performed with EZ DNA Methylation-Gold (Zymo Research) according to the manufacturer’s instructions. The final amplification was performed with KAPA HiFi U+ (Roche Sequencing), IDT xGen Primers (20 μM - Integrated DNA Technologies), and 12 PCR cycles.

Library QC was performed using the D1000 Assay on a 4200 Agilent TapeStation, and its concentration measured with the Qubit dsDNA BR Assay (LifeTechnologies). Libraries were sequenced on a NovaSeq6000 (S4 lane) as paired-end 150 bases.

### Data Processing

Demultiplexed Fastq files were subject to QC (FastQC - Babraham Bioinformatics) and trimming with cutadapt v2.10 (Martin 2011) (options: -u −10 -U 10 -q 20 -m 50) before alignment to the *Xenopus tropicalis* genome (version XENTR_10.0) with BSBolt Align (Farrell et al. 2021) (default options). PCR duplicates were removed with samtools markdup (option -r) v1.15 (Danecek et al. 2021). DNA methylation was called using BSBolt CallMethylation v1.3.0 (options: -BQ 10 -MQ 20 -IO) resulting in CGmap files. The matrices of common CpG sites (with at least 3x or 5x coverage) were produced using BSBolt AggregateMatrix. Data for metagene plots were calculated with CGmap tools (tools: bed2fragreg; mfg) (Guo et al. 2018). Data for chromosome-wide DNA methylation distribution were calculated with CGmap tools (tool: mbin).

Data from human mammary epithelial tissues were downloaded from the Encode database (doi:10.17989/ENCSR656TQD, file ENCFF699GKH).

All plots were generated in R (version 4.1.2). PCA was performed with the PCAtool package (v2.6.0) in R (Blighe K and Lun A 2022). The methylation clock model was created with the ElasticNet function from the Python module sklearn.linear_model (Pedegrosa et al., 2011). The chosen hyperparameters (alpha=0.00283693 and l1_ratio=0.5) were found via grid search with a LOOCV procedure with the ElasticNetCV function. The correlation and ANOVA analyses to select sites associated with the three different age groups were performed using the cor.test (Pearson) and anova functions in R, respectively. The top 4500 sites (ranked by correlation and ANOVA adjusted p-values) were selected for probe design, resulting in a final design capturing approximately 3000 sites of the initial list.

## Supplementary Materials

**Supplementary Figure 1:**
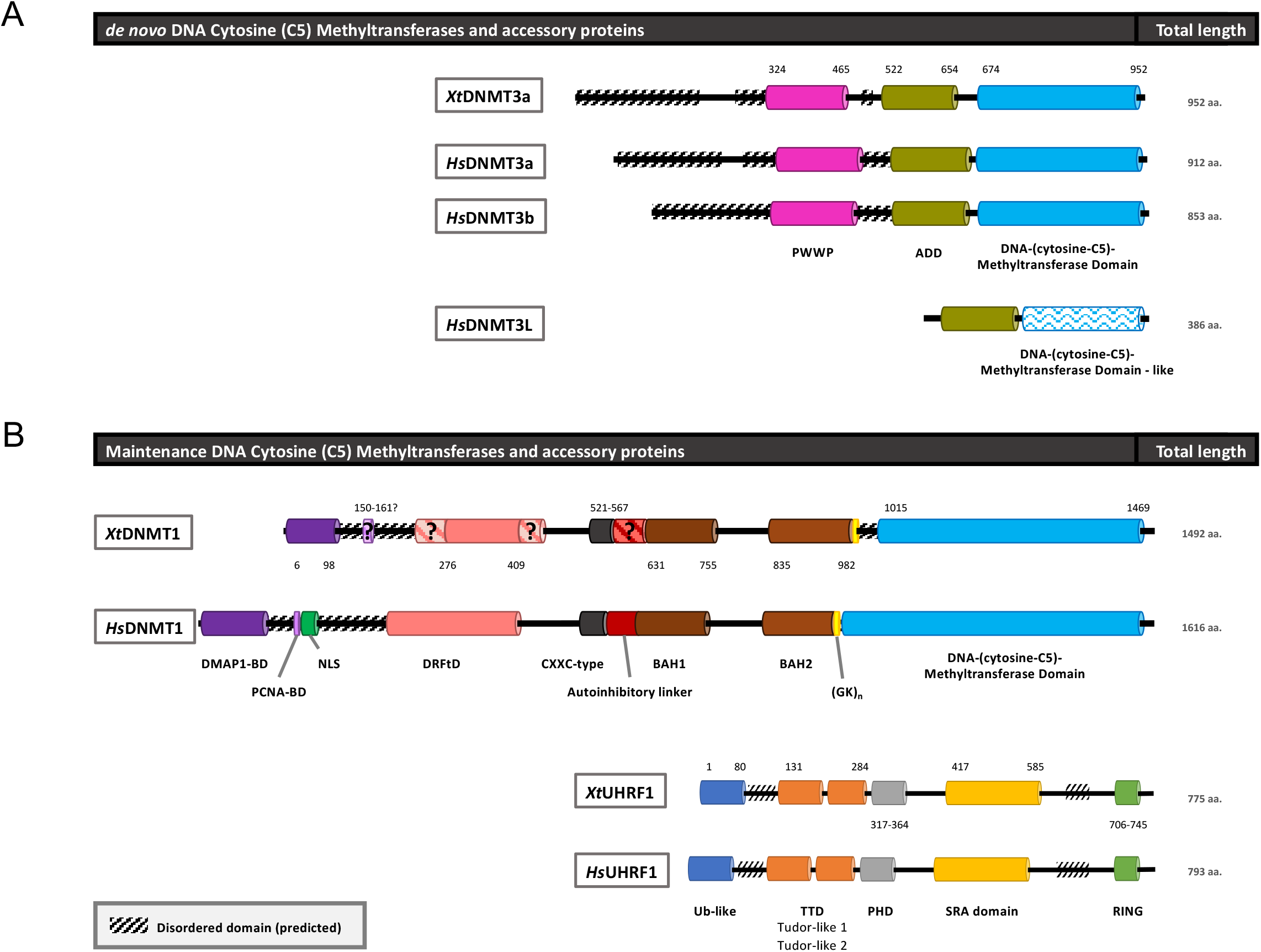
Comparison of DNA methyltransferases and main accessory proteins from H. sapiens and X. tropicalis. (A) De novo DNA methyltransferases: XtDNMT3a (accession number F7BU15), HsDNMT3a (Q9Y6K1), HsDNMT3b (Q9UBC3), and HsDNMT3L (Q9UJW3). PWWP domain: Pro-Trp-Trp-Pro motif (fuchsia); ADD domain: ATRX-DNMT3-DNMT3L, comprised of an N-terminal C2C2-type zinc finger (GATA-like), an imperfect PHD-type C4C4 zinc finger PHD finger and a C-terminal α-helix (olive). DNA-(cytosine-C5)-Methyltransferase Domain: catalytic domain (light blue). DNA-(cytosine-C5)-Methyltransferase Domain – like: inactive methyltransferase domain in DNMT3L (white with light blue zig-zags). (B) Maintenance DNA methyltransferases: XtDNMT1 (F6QE78); HsDNMT1 (P26358); XtUHRF1 (F6UA42); HsUHRF1 (Q96T88). DMAP1-BD: interaction module that binds the transcriptional co-repressor DMAP1 (DNA methyltransferase-associated protein 1) (purple). PCNA-BD: Proliferating cell nuclear antigen interaction domain (lavender). In XtDNMT31, the PCNA-BD is colored with lavender stripes because no domain was predicted in this region despite the high levels of sequence identity in this region. NLS: nuclear localization signal (green). No prediction present for XtDNMT1. DRFtD: DNA replication foci-targeting domain (salmon). Despite the sequence similarity, the predicted domain in XtDNMT1 is shorter than the human counterpart (salmon diagonal stripes in the missing regions). CXXC-type: CXXC zinc finger domain (dark grey). Autoinhibitory linker, not predicted in XtDNMT1 despite high sequence similarity (red/diagonal red stripes). BAH1/BAH2: Bromo-adjacent homology domain (brown/dark orange). (GK)n: glycine-lysine (GK) repeats (bright yellow). DNA-(cytosine-C5)-Methyltransferase Domain: catalytic domain (light blue). Ub-like: ubiquitin-like domain (blue). TTD: tandem Tudor domain (orange). PHD: Plant Homeo Domain finger (grey). SRA: SET and RING-finger Associated domain containing the YDG motif, 5meC binding pocket, and residues important for base flipping (gold). RING: RING (really interesting new gene) zinc-finger (light green). Diagonal black-striped boxes indicate a predicted disordered consensus sequence. Domains in the Xt proteins have been compiled from InterProScan predictions.

**Supplementary Figure 2:**
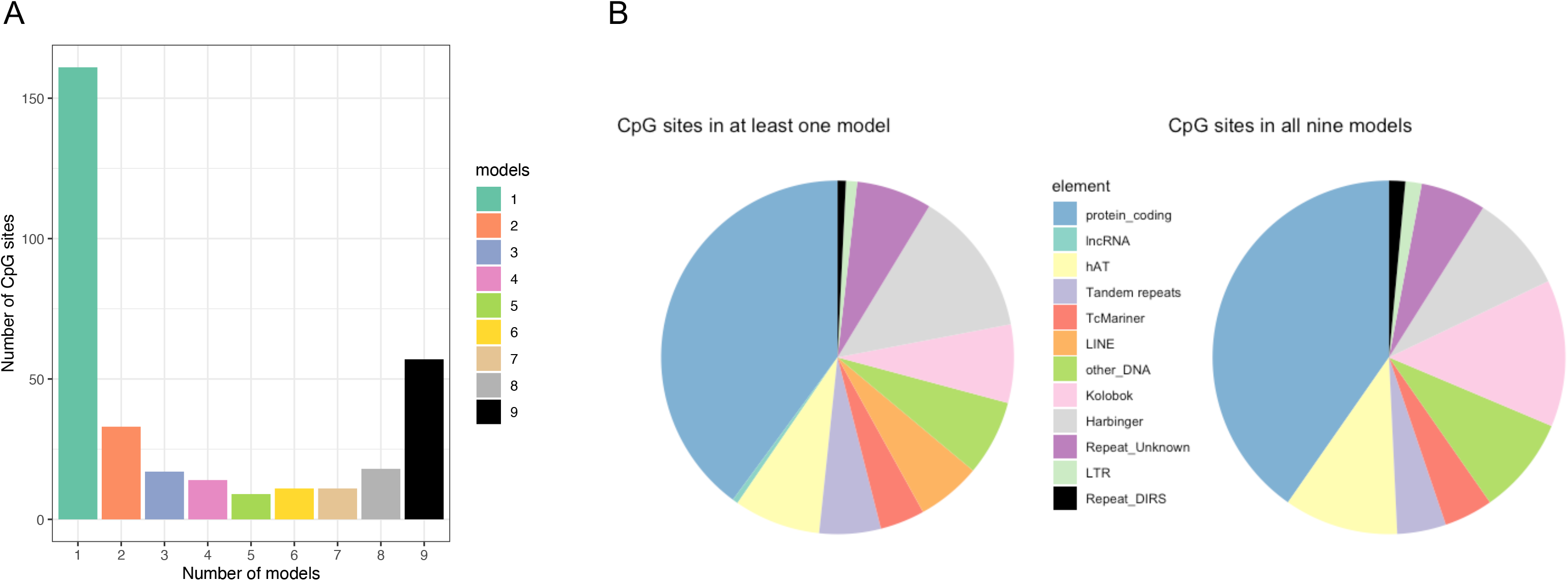
Predictive CpG sites. (A) Number of CpG sites used in N models. E.g. The CpG sites in the category “Number of models = 9” correspond to the sites employed by all 9 models; the CpG sites in the category “Number of models = 5” correspond to the sites employed in 5 models only (9 models total). (B) Distribution of the closest genomic elements for all CpG sites used in at least one model (left) and for the CpG sites common to all 9 models (right).

**Supplementary Figure 3:**
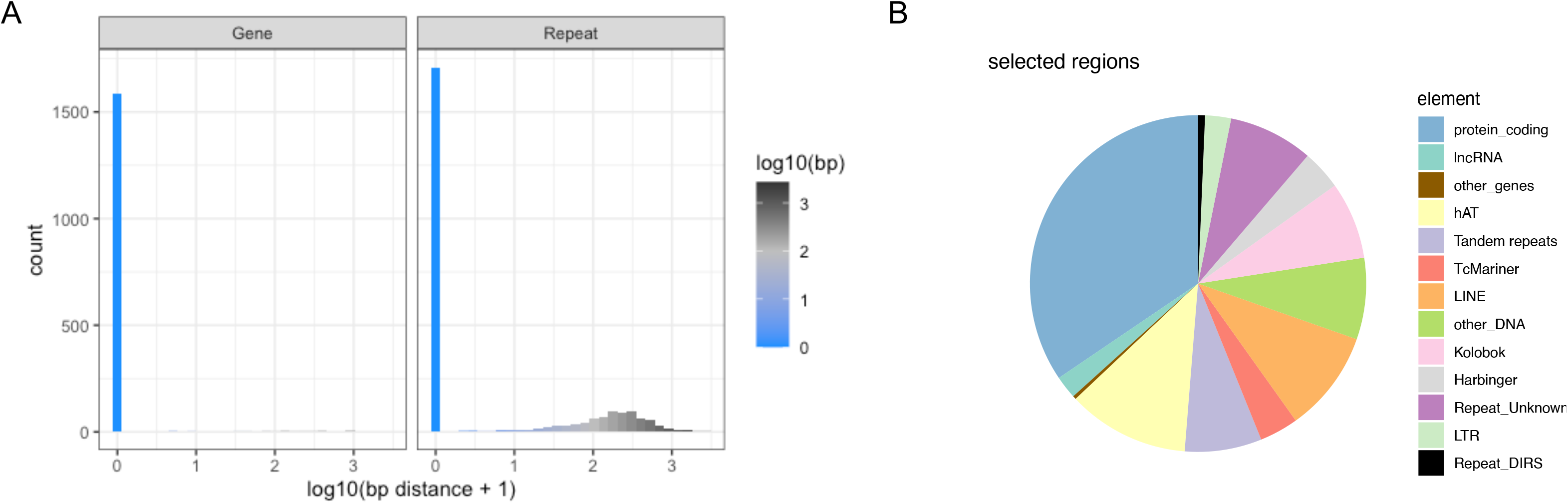
(A) Distribution of distances between selected regions and genome elements (genes and repeats). (B) Distribution of the closest genomic elements (both genes and repeats) for each selected region.

